## Supplementary Data for "Assessment of Immunogenicity and Efficacy of CV0501 mRNA-based Omicron COVID-19 Vaccination in Small Animal Models"

#### Supplementary Figure 1: CV0501 vaccine induced strong cross-nAbs at Day 42 post-immunization

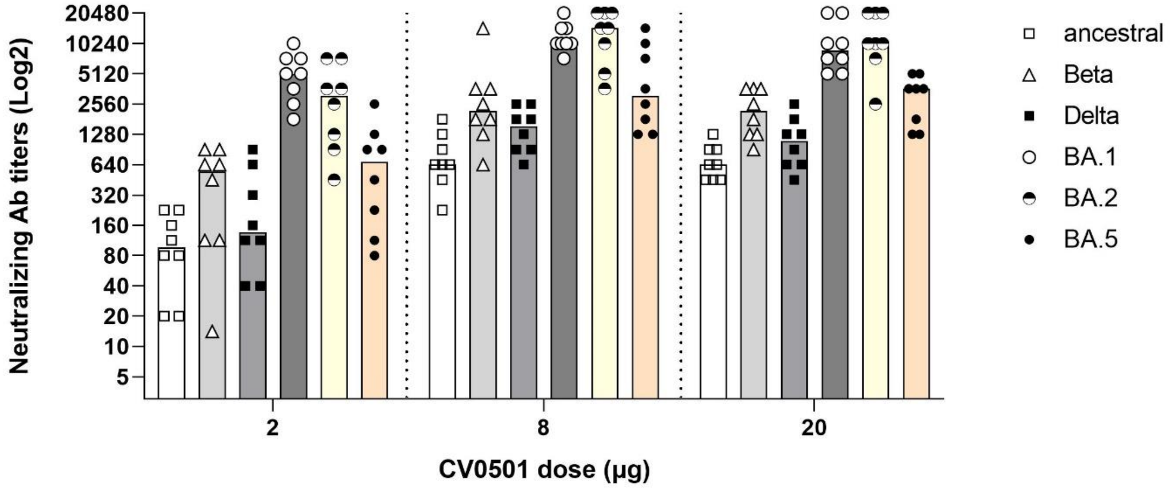


**Figure S1: CV0501 vaccine induced strong cross-nAbs at Day 42 post-immunization** (Data from Fig 1 and 2 displayed).

Wistar rats (n=8/group) were immunized i.m. on Days 0 and 21 with 2, 8 or 20 µg CV0501. nAbs against ancestral, Beta, Delta, BA.2 and BA.5 SARS-CoV-2 were assessed from serum isolated on Day 42 post-immunization. Each symbol represents an individual animal and bars depict the median value.

#### Supplementary Figure 2: CV0501 induces dose dependent IFN-γ secreting T cells

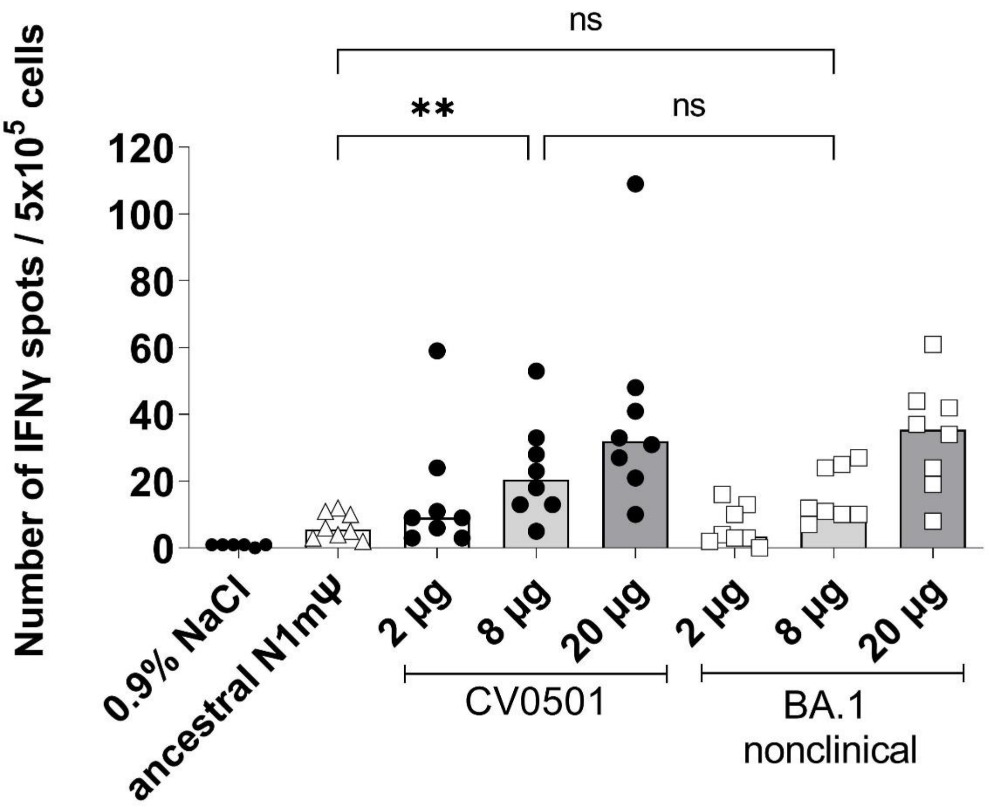


**Figure S2: Immunization with CV0501 induced significantly higher IFN-γ secreting T cells compared with ancestral N1mΨ.**

Splenocytes from rats immunized on Days 0 and 21 with either CV0501 (2, 8 or 20 µg), BA.1 nonclinical (2, 8 or 20 µg), ancestral N1mΨ (8 µg), or 0.9% NaCl buffer (sham control) were isolated on Day 42 and single-cell suspensions were prepared and stimulated for 24 hours at 37°C using a SARS-CoV-2 Omicron peptide library (JPT, PM-SARS2-SMUT08-1) at 1 µg/mL. Each dot represents an individual animal and bars depict the mean. Statistical analysis was performed using ANOVA and Dunn's multiple comparison test (**; $p=0.0049$).

#### Supplementary Figure 3: CV0501 booster immunization induces high cross-neutralizing antibodies against BA.1 in Wistar rats

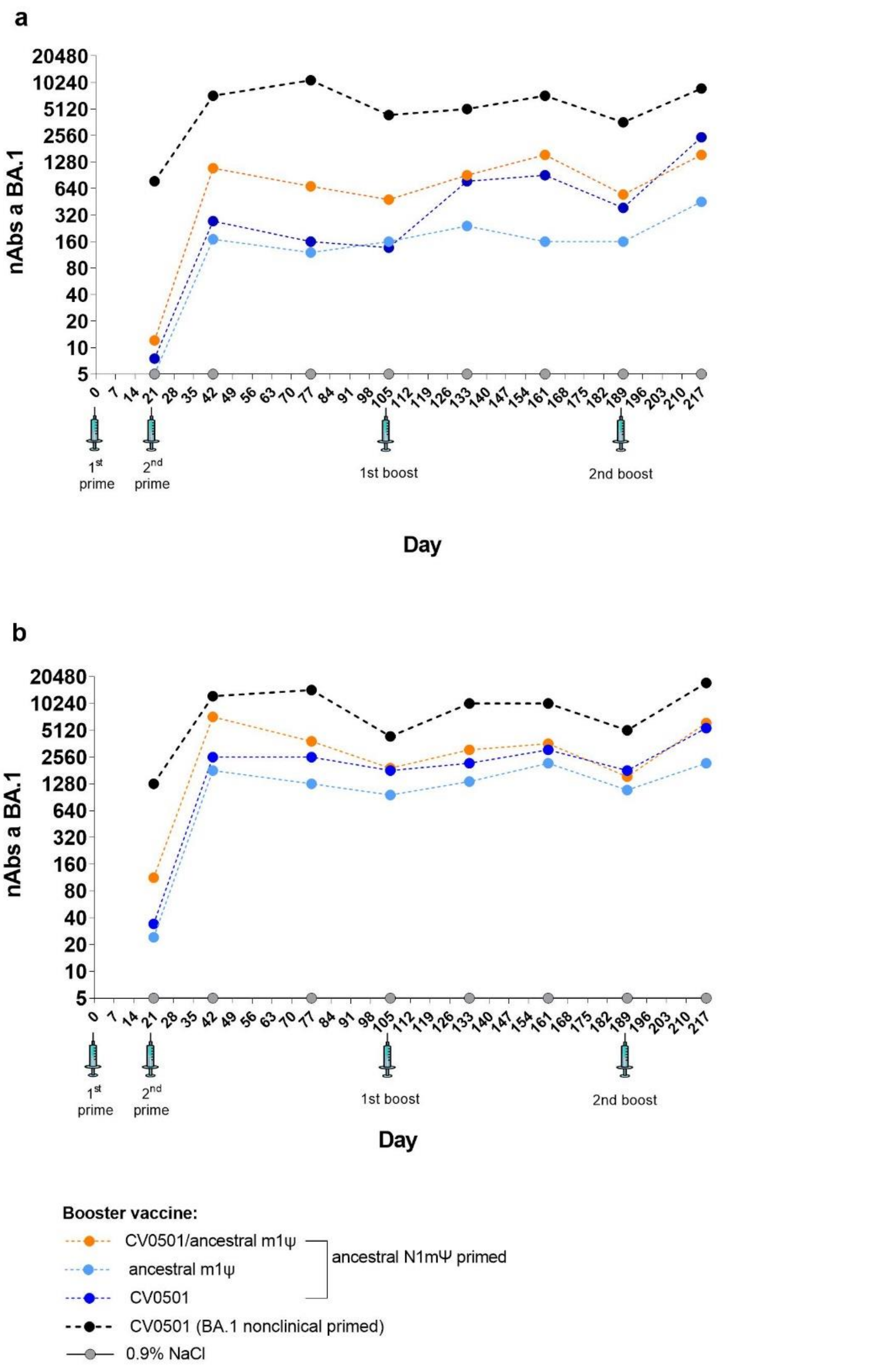

**Figure S3: CV0501 booster immunization induces high cross-neutralizing antibodies against BA.1 in Wistar rats**

Wister rats (n=8/group) were immunized on Days 0 and 21 with either 2 µg (**a**) or 8 µg (**b**) ancestral N1mΨ or BA.1 nonclinical. On Days 105 and 189, rats were given third and fourth (booster) doses of either CV0501 alone, ancestral N1mΨ alone, or bivalent CV0501/ancestral N1mΨ (half doses of each) at 2 µg or 8 µg doses. 0.9% NaCl was used as a sham control and was administered at the same timepoints. nAbs against BA.1 SARS-CoV-2 were assessed in serum obtained on Days 21, 42, 77, 105, 133, 161, 189 and 217 and analyzed at each timepoint. Each symbol represents the median value.

#### Supplementary Figure 4: Induction of nAb against BA.1 Omicron variants observed after CV0501 immunization.

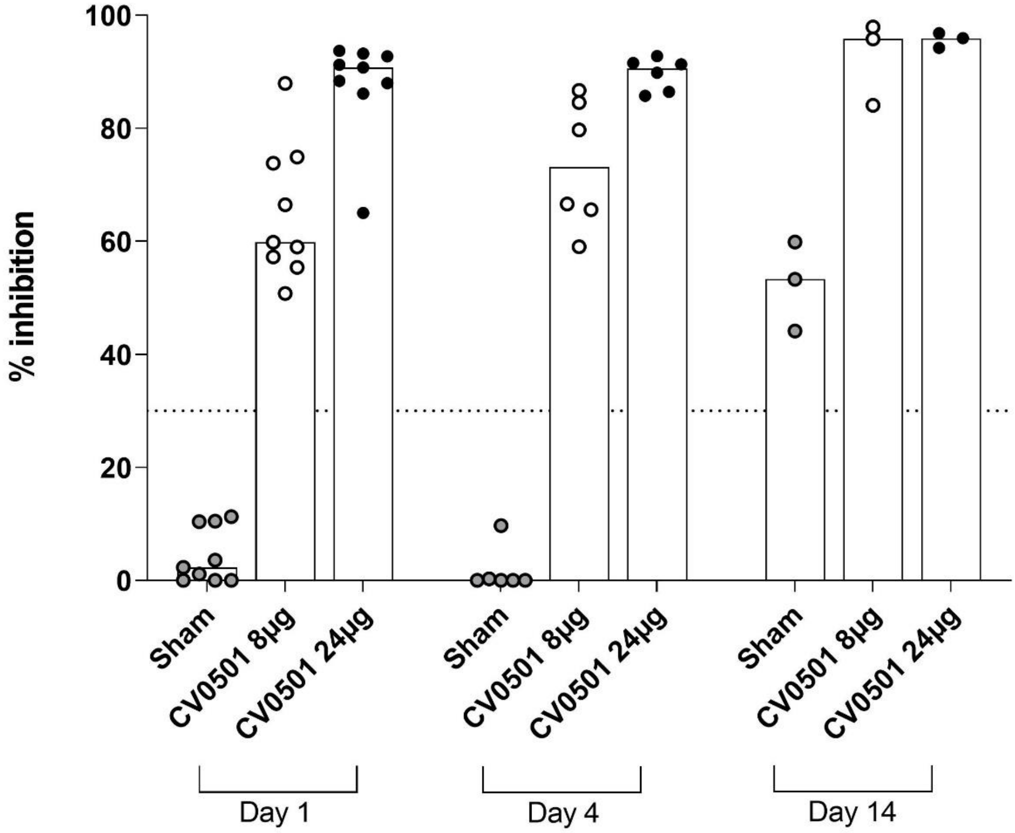


**Figure S4: Percentage of inhibition against BA.1 Omicron variants observed after CV0501 immunization.**

Hamsters (n=9/group) were immunized on Days 0 and 28 with 8 µg or 24 µg doses of CV0501 or 0.9% NaCl (sham controls) and challenged on Day 56 with Omicron BA.2 ($10^5$ $TCID_{50}$/animal administered i.n. at 0.05 mL per nostril). A surrogate ELISA was performed using serum samples to assess the percentage of inhibition against BA.1 on Days 0 and 4 (6 animals/groups) or Day 14 (3 animals/groups) post-challenge. Each dot represents an individual animal, bars depict the median value of the group at the various time points. Horizontal line represents the threshold of 30% inhibition, above which animals are classified as sero-positive against BA.1.

### Supplementary Methodology

#### Supplementary Table 1: Experimental design for booster response of monovalent and bivalent CV0501

| Group | Animals | 1st prime (Day 1)<br>2nd prime (Day 21) | 1st boost<br>(Day 105) | 2nd boost<br>(Day 189) | Blood collection schedule |
|---|---|---|---|---|---|
| 1 | Female Wistar rats (N=6) | 0.9% NaCl | 0.9% NaCl | 0.9% NaCl | Days 21, 42, 77, 105, 133, 161, 189, and 217 |
| 2 | Female Wistar rats (N=8) | ancestral N1mΨ 2 µg | ancestral N1mΨ 2 µg | ancestral N1mΨ 2 µg | |
| 3 | | ancestral N1mΨ 8 µg | ancestral N1mΨ 8 µg | ancestral N1mΨ 8 µg | |
| 4 | | ancestral N1mΨ 2 µg | CV0501 2 µg | CV0501 2 µg | |
| 5 | | ancestral N1mΨ 8 µg | CV0501 8 µg | CV0501 8 µg | |
| 6 | | ancestral N1mΨ 2 µg | Bivalent CV0501/ancestral N1mΨ2 µg (total) | Bivalent CV0501/ancestral N1mΨ 2 µg (total) | |
| 7 | | ancestral N1mΨ 8 µg | Bivalent CV0501/ancestral N1mΨ 8 µg (total) | Bivalent CV0501/ancestral N1mΨ 8 µg (total) | |
| 8 | | BA.1 nonclinical N1mΨ 2 µg | CV0501 2 µg | CV0501 2 µg | |
| 9 | | BA.1 nonclinical N1mΨ 8 µg | CV0501 8 µg | CV0501 8 µg | |

N1mΨ, N1-methylpseudouridine.

#### Supplementary Table 2: Antigens included in ACE2 inhibition assay

| SARS-CoV-2 variant | Manufacturer | Cat. # | Lot # |
|---|---|---|---|
| Wild type (B1 isolate) | NMI | – | – |
| Alpha | NMI | – | – |
| Beta | NMI | – | – |
| Gamma | NMI | – | – |
| Delta | NMI | – | – |
| Omicron BA1 (15 RBD mutations) | Sino Biological | 40592-V08H121 | LC16FE2107 |
| Omicron BA1 (12 RBD mutations) | Sino Biological | 40592-V08H122 | LC16MA2204 |
| Omicron BA2 | Sino Biological | 40592-V08H123 | MA16MA1803 |
| Omicron BA2.12.1 | Sino Biological | 40592-V08H132 | MB16MY1181 |
| Omicron BA4 | Sino Biological | 40592-V08H130 | LC16JU1717 |
| Omicron BA5 | Sino Biological | 40592-V08H131 | MA16JL0542 |

RBD, receptor binding domain; SARS-CoV-2, severe acute respiratory syndrome coronavirus-2.
